# Efficient generation of human dendritic cells from iPSC by introducing a feeder-free expansion step for hematopoietic progenitors

**DOI:** 10.1101/2024.06.04.594010

**Authors:** Zahra Elahi, Vanta Jameson, Magdaline Sakkas, Suzanne K Butcher, Justine D Mintern, Kristen J Radford, Christine A Wells

## Abstract

Dendritic cells (DCs) are rare innate immune cells that are essential regulators of anti-tumour, anti-viral and vaccine responses by the adaptive immune system. Conventional dendritic cells, particularly the cDC1 subset, are most desired for DC-based immunotherapies, however, it can be difficult to isolate sufficient numbers of primary cells from patients. The most common alternate sources of DC are *ex vivo*, such as monocyte-derived or DC expanded from cord blood hematopoietic progenitors. Induced pluripotent stem cells (iPSC) offer a promising solution, providing an opportunity for *in vitro* generating DCs that are suitable for patient-derived or off-the-shelf batch-manufactured cells. Here, we developed an *in vitro* protocol designed to maximise the yield of iPSC-derived DC progenitors, with the specific goal of generating DC1-like cells. The iPSC-DCs subsets generated by our method could be partitioned by cell surface phenotypes of cDC1, cDC2 and DC3, but they were most transcriptionally similar to monocyte-derived DC (MoDC). Stimulated iPSC-DCs generated pro-inflammatory cytokines, expressed migratory chemokine receptors including CCR7 which indicates capacity to traffic through lymphatic endothelium, and upregulated co-stimulatory molecules, indicating their potential for productive interactions with T-cells. This method offers a promising step towards an expandable source of allogeneic human dendritic cells for future applications.

## Introduction

Dendritic cells (DC) are specialised antigen-presenting cells that traffic to lymph nodes where they efficiently activate naïve CD4+ and CD8+ T lymphocytes. They can perform both conventional presentation and cross-presentation (presenting exogenous antigens through MHC-I molecules) of antigens to T cells and therefore are the target cell types for most vaccines (1). DC-based vaccines have shown sporadic but highly effective potential in anti-cancer immunotherapies for late-stage melanoma (2), glioma (3) and other aggressive tumours (reviewed by Wculek et al. (4)).

Among the heterogenous population of DCs, the cDC1 subset has an exceptional ability to cross-present tumour antigens to CD8+ T cells and induce an efficient antigen-specific response (5, 6). Murine cDC1 displayed a higher capacity to activate and induce the proliferation of CD8+ T cells than cDC2 and MoDC subsets (7) and were shown to be the only antigen-presenting cells able to transport intact tumour antigens to the tumour-draining lymph nodes and prime CD8+ T cells (8). However, the clinical application of human DCs generally, and cDC1 subsets in particular, is fundamentally limited by their scarcity in circulation, short half-life and poor proliferative abilities. Together, these prevent efficient manufacturing of primary cDC1 at scale.

Currently, the most common approach for producing clinical-grade DCs is the *ex vivo* differentiation of monocytes to dendritic cells. To obtain Monocyte-derived DC (MoDC), the CD14+ fraction of peripheral blood mononuclear cells (PBMC) are isolated and treated with IL4 and GM-CSF for several days (9). MoDCs show a typical dendritic morphology and express MHC class I and class II molecules, CD172a, CD1a, CD1c and CD11c (reviewed by (10)) with variable CD14 expression (11, 12). MoDCs matured with LPS and TNFα have enhanced expression of MHC II and co-stimulatory molecules and generate high levels of proinflammatory cytokines with the capacity to induce naïve CD4+ T cell differentiation (13). MoDCs have been the cell of choice for many clinical studies using autologous DC-based vaccines (14). However, monocytes and MoDC are short-lived cells (15, 16), and the number of MoDC available is fundamentally constrained by the number of monocytes that can be isolated from a donor, impacting their manufacturing or delivery for clinical use.

An alternate source is the differentiation of CD34+ hematopoietic stem cells isolated from cord blood (CB-HSC). These cultures are able to support the differentiation of multiple DC subsets, such as CLEC9A+ cDC1 (17, 18), CD123+ pDC, CD1c+ cDC2 and MoDC (19, 20). Nevertheless, one of the challenges associated with the generation of DC, particularly cDC1 subset, from CB-HSC is the low yield of differentiation (17, 21, 22). Transcriptionally, we have shown that CB-DC captures an immunoregulatory profile (mregDC) (22), which leads to less efficiency in anti-cancer responses due to the induction of many regulatory genes (23). While the phenotype of these cells partially matches the primary cDC1 or cDC2, manufacturing of CB-cDC remains a major limitation when seeking to generate sufficient DCs for clinical application.

Pluripotent stem cells offer another alternative for DC derivation, and two broad approaches have been reported. The first approach which is more common uses embryoid bodies (EB) to produce HSC followed by differentiation toward terminal DCs in a feeder-free culture setup. The absence of feeder cells in this method makes it a well-defined process; however, it generated monocyte- and cDC2-like cells from iPSC (24–26). The second approach generated conventional DCs in a co-culture of iPSC with mouse stromal cells engineered to express Notch ligand DLL1 (27). Although this approach derives a high percentage of cDC1, using mouse feeder cells introduces xenogeneic and pathogenic components to the system, which makes their clinical applications challenging. Therefore, we designed a feeder-free EB-based method to generate and expand iPSC-derived HSC that potentially differentiate into multiple iPSC-DC subsets with proinflammatory functions.

## Methods

### Cell lines and ethics

All works conducted on primary human cells, and iPSC lines was overseen by the University of Melbourne HREC under approvals 1646608 and 2023-28021-46872-3.

Primary cord blood cells were obtained through the BDMI cord blood bank at the Royal Children Hospital, Melbourne.

This study also used two human iPSC cell lines previously described (28) including PB001.1 (hPSC reg: MCRIi001-A; RRID:CVCL_UK82) (29), obtained from the Stem Cell Core Facility at the Murdoch Children’s Research Institute and HDF51(30) (RRID:CVCL_UF42), which was a gift from Professor Andrew Laslett, CSIRO.

### iPSC maintenance and expansion

iPSCs were cultured in supplemented E8 media (Thermo Scientific, #A1517001) on Matrigel® Matrix (Thermo Scientific, #354277) coated dishes. To coat the dishes, 1 ml of frozen Matrigel aliquot was thawed at 4°C overnight. Thawed Matrigel was diluted 1/50 in cold E8 media and was then added to the cell culture dishes (Merck, #Z755923). Coated dishes were incubated at room temperature for 1 hour and used immediately or kept in the fridge for later application.

Seeded iPSC cells were maintained in a humidified incubator at 37°C with 5% CO2 and spent media was replaced with fresh supplemented E8 every day. To passage cells when confluent at 70-80%, cells were first washed with PBS (Thermo Scientific, #10010023), then incubated with 1/1000 EDTA (Thermo Scientific, #15575020) in PBS buffer of for 3-4 minutes at 37°C with 5% CO2 to allow detachment. After incubation, the buffer was gently removed, and cells were detached by pipetting using fresh E8 media. Detached cells were collected in a 15ml centrifuge tube (Corning, #430791), spun with centrifuge (Techcomp, #CT15RT) at 300g for 5min and reseeded at ¼ dilution.

### EB formation and differentiation

EB culture medium was based on STAPEL media (31, 32) (Table 1). To generate EBs, iPSC cells were detached enzymatically using cell release buffer (1/1000 EDTA/PBS), and clumps of cells were collected by pipetting using EB media (Table 1) supplemented with differentiation factors for day 0-1 (Table 2). Collected cell clumps were filtered through 70 µm-sized strainers into a 50ml sterile tube. Cell clumps were removed by 10ml serological pipettes and added gently to ultra-low attachment culture dish (Sigma Aldrich, cat#CLS3261) to be maintained at 37°C with 5% CO2 placed on an orbital shaker (SCIENTIFIX, cat#NBT-101SRC) rotating at 32rpm. For each 100mm dishes, 2 to 3×10^6^ cells were used for EB formation. Media change strategy, including the details of each differentiation factor, is described in Table 2. To improve cell viability at the EB formation step, 0.2 nM ROCK Inhibitor Y-27632 (Stemcell Technologies, #72307) was added to the media day 0-1 and discontinued since then. From Day 7, iPSC-derived hematopoietic cells started to emerge from embryoid bodies as suspension cells.

**Table 1.** Components of EB culture media.

| Components | Supplier/ catalogue number | Stock conc. | Final conc. | Volume for 50ml media |
| --- | --- | --- | --- | --- |
| Iscove's Modified Dulbecco's Medium; no phenol red (IMDM) | Thermo Fisher/ 21056023 | - | - | 33.25 |
| F-12 nutrient mixture with Glutamaxl (F12) | Thermo Fisher/ 31765035 | - | - | 12.5 |
| Protein-free hybridoma mix (PFHMII) | Thermo Fisher/ 12040077 | - | - | 2 |
| BovoStar Bovine Serum Albumin (Australian origin) (BSA) | Bovogen biologicals/ BSASAU | 10% | 0.25% | 1.25 |
| Poly vinyl alcohol (PVA) | Sigma-Aldrich/ P8136 | 10% | 0.15% | 750 |
| Methylcellulose | Sigma-Aldrich/ M6385 | 10% | 0.10% | 500 |
| Insulin-Transferrin-Selenium-Ethanolamine (ITS-X) |  | 100% | 1x | 500 |
| L-Ascorbic acid 2-phosphate (AA2P) | Sigma-Aldrich/ A8960 | 5 mg/ml | 50 µg/ml | 500 |
| GlutaMAX | Thermo Fisher/ 35050061 | 200 mM | 2 mM | 500 |
| Linoleic and Linolenic acids (L+L) | Sigma-Aldrich/ Linoleic acid (L1012), Linolenic acid (L2376) | 5000x | 1x | 10 |
| Synthetic cholesterol | Sigma-Aldrich/ M6385-250G | 20 mg/ml | 2.2 µg/ml | 5.5 |
| 2-Mercaptoethanol (B-ME) | Sigma-Aldrich/ M3148 | 14.3M | 0.11 nM | 0.38 |

**Table 2.** Media change strategy of EB differentiation. rh stands for Recombinant Human proteins.

| Components | Supplier/ catalogue number | Concentration of factor in EB media in each media change |  |  |  |  |
| --- | --- | --- | --- | --- | --- | --- |
|  |  | Day 0-1 | Day 1-2 | Day 2-4 | Day 4-7 | Day 7-onward |
| **rh Activin A** | R&D Systems/ 338AC010 | 5 ng/ml | - | - | - | - |
| **rh CHIR** | R&D Systems/ 442310 | 0.2 μM | 0.2 μM | - | - | - |
| **rh BMP-4** | R&D Systems/ 314BP050 | 20 ng/ml | 20 ng/ml | 20 ng/ml | 20 ng/ml | - |
| **rh FGF** | PeptoTech/ AF-100-18B | 5 ng/ml | 10 ng/ml | 10 ng/ml | 10 ng/ml | 10 ng/ml |
| **rh VEGF** | PeptoTech/ AF-100-20 | 40 ng/ml | 40 ng/ml | 40 ng/ml | 40 ng/ml | 20 ng/ml |
| **rh SCF** | PeptoTech/ AF-300-07 | 40 ng/ml | 40 ng/ml | 40 ng/ml | 40 ng/ml | 20 ng/ml |
| **rh FLT3-L** | PeptoTech/ AF-300-19 | - | - | - | 50 ng/ml | 50 ng/ml |
| **rh IL-3** | PeptoTech/ AF-200-03 | - | - | - | 25 ng/ml | 25 ng/ml |

### iPSC-derived progenitor expansion

Progenitor cells that emerged from EBs were collected and spun at 300g for 5min. Collected cells were resuspended in 1ml of pre-warmed Amplification media described in Table 3. Cells were cultured in 100,000 cell/ml density on T25 cell culture flasks and incubated at 37°C with 5% CO2. Cells were expanded 4-7 days based on the experimental design. During expansion media was not changed.

**Table 3.**
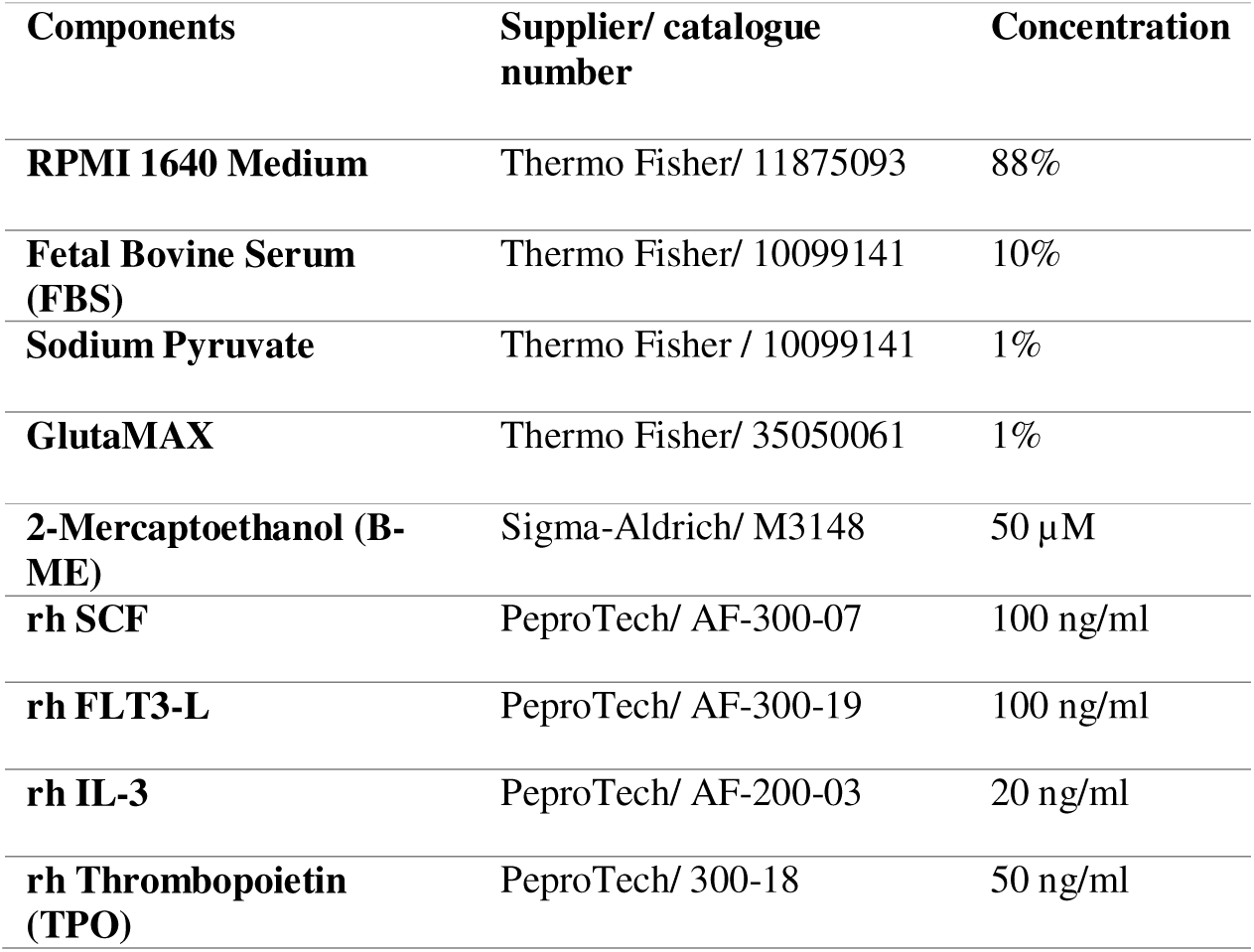
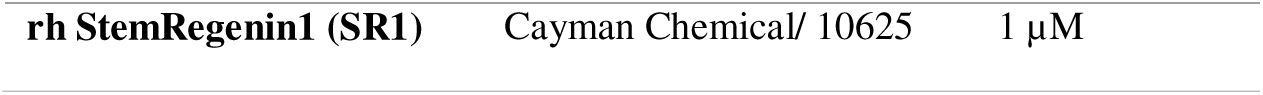
Components of Amplification media.

| Components | Supplier/ catalogue number | Concentration |
| --- | --- | --- |
| **RPMI 1640 Medium** | Thermo Fisher/ 11875093 | 88% |
| **Fetal Bovine Serum (FBS)** | Thermo Fisher/ 10099141 | 10% |
| **Sodium Pyruvate** | Thermo Fisher / 10099141 | 1% |
| **GlutaMAX** | Thermo Fisher/ 35050061 | 1% |
| **2-Mercaptoethanol (B-ME)** | Sigma-Aldrich/ M3148 | 50 μM |
| **rh SCF** | PeptoTech/ AF-300-07 | 100 ng/ml |
| **rh FLT3-L** | PeptoTech/ AF-300-19 | 100 ng/ml |
| **rh IL-3** | PeptoTech/ AF-200-03 | 20 ng/ml |
| **rh Thrombopoietin (TPO)** | PeptoTech/ 300-18 | 50 ng/ml |
| **rh StemRegenin1 (SR1)** | Cayman Chemical/ 10625 | 1 μM |

### iPSC-derived Dendritic Cell differentiation

Expanded progenitors were collected and spun at 300g for 5min. 200,000 cell/ml were seeded on T25 tissue culture flasks in DC differentiation media described in Table 4. At day 6, half of media was replaced with fresh differentiation media with 2X concentration of cytokines.

**Table 4.**
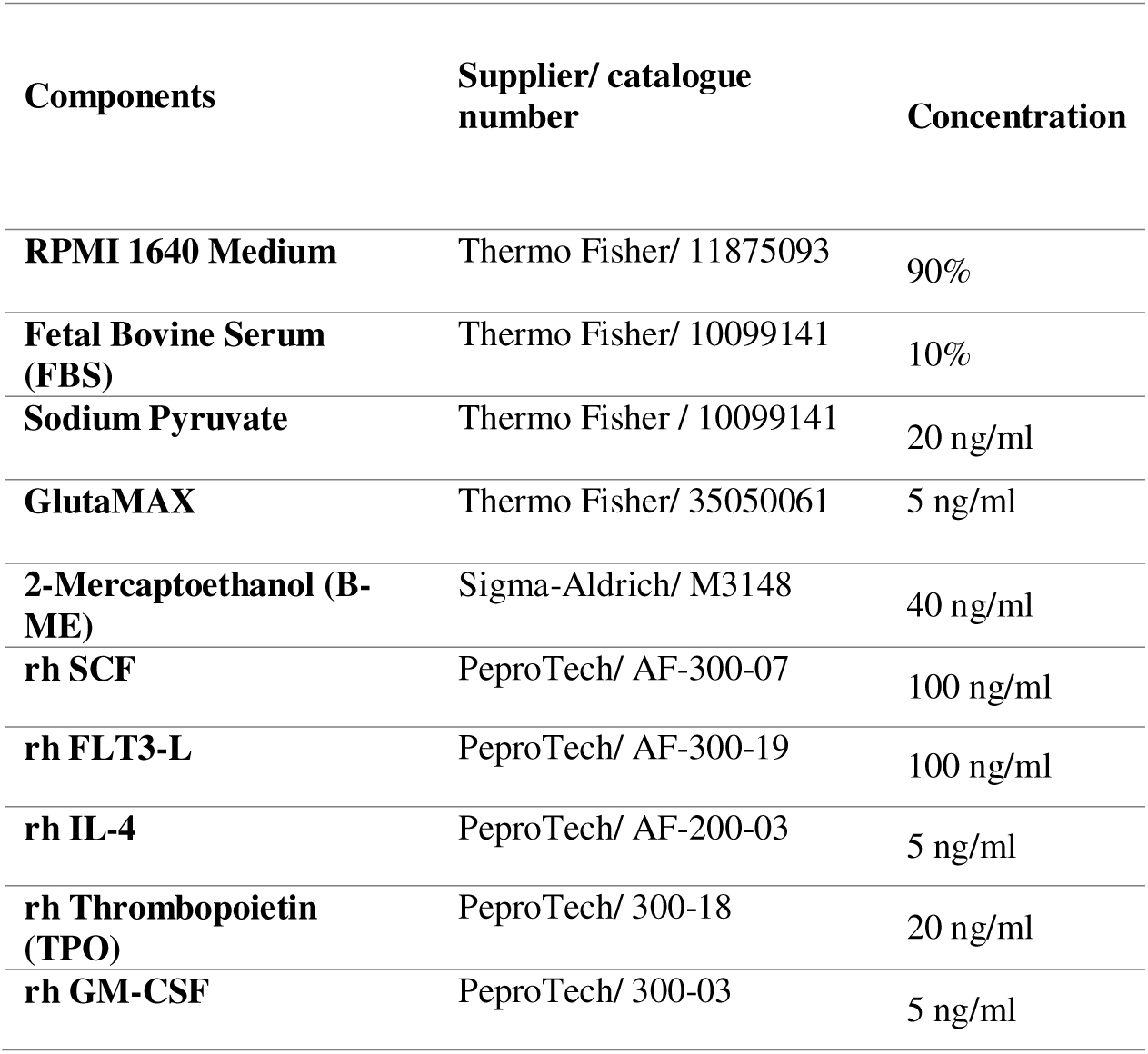
Components of DC differentiation media.

### Cord blood CD34+ HSC-derived DC differentiation

The CD34+ HSC cells were isolated using Human Cord Blood CD34 Positive Selection Kit II (Stemcell Technologies, #17896) as per the supplier instruction. Briefly, first step of enrichment used RosetteSep cocktail to deplete platelets. Then CB were diluted 1:1 (v/v) into a dilution buffer (1mM EDTA in PBS supplemented by 2% FBS (Thermo Fisher, #10099141)) and undelayed with density gradient medium LymphoPrep (Stemcell Technologies, #07801). The mix were centrifuged at 1200×g for 20 mins with no brake. Blood mononuclear cells (BMC) were harvested from the interface layer between plasma and red cell and spun at 300×g for 10 mins with no brake. The cell pellet diluted 1:1 (v/v) in DB and transferred to a 5ml round bottom polystyrene tube (Stemcell Technologies, #38030). CD34+ cells were enriched by magnetic selection using Selection antibody cocktail in the kit. Isolated cells were plated in amplification media Table 5 for 4-7 days. Expanded cells were harvested and used for characterisation, freezing or proceeding to the differentiation phase. For the latter purpose, cells were cultured in DC differentiation media (Table 6) for 11 days; at day 6, half of the media was replaced with fresh media containing 2X concentration of cytokines.

**Table 5.** List of flow cytometry antibodies and Isotype controls.

| Antibodies or Isotypes | Supplier | Catalogue number | Dilution |
| --- | --- | --- | --- |
| PE anti-human CD370 (CLEC9A/DNGR1) | BioLegend | 353804 | 1/50 |
| PE/Dazzle™ 594 anti-human CD1c | BioLegend | 331532 | 1/200 |
| Alexa Fluor® 647 anti-human CD141 (Thrombomodulin) | BioLegend | 344124 | 1/50 |
| PE/Cyanine7 anti-human CD11c Antibody | BioLegend | 337216 | 1/200 |
| Alexa Fluor® 700 anti-human CD172a/b (SIRPα/β) | BioLegend | 323815 | 1/100 |
| Brilliant Violet 421™ anti-human CD14 | BioLegend | 325628 | 1/100 |
| PE/Dazzle594 anti-human CD135 (Flt-3/Flk-2) | BioLegend | 313320 | 1/50 |
| Alexa Fluor® 488 anti-human HLA-DR | BioLegend | 307619 | 1/100 |
| PE/Cyanine7 anti-human CD115 (CSF-1R) | BioLegend | 347308 | 1/100 |
| APC anti-human CD116 | BioLegend | 305914 | 1/200 |
| PE anti-human CD131 | BioLegend | 306104 | 1/200 |
| PE anti-human CD34 | BioLegend | 343506 | 1/200 |
| APC anti-human CD45 | BioLegend | 304012 | 1/200 |
| APC/Cyanine7 anti-human CD40 | BioLegend | 334323 | 1/100 |
| PE anti-human CD80 | BioLegend | 374205 | 1/100 |
| Brilliant Violet 421™ anti-human CD86 | BioLegend | 305425 | 1/100 |
| PE Mouse IgG1, κ Isotype | BioLegend | 400112 | 1/50 |
| APC Mouse IgG1, $\kappa$ Isotype | BioLegend | 400120 | 1/100 |
| PE/Dazzle™ 594 Mouse IgG1, $\kappa$ Isotype | BioLegend | 400177 | 1/200 |
| Alexa Fluor® 647 Mouse IgG1, $\kappa$ Isotype | BioLegend | 400130 | 1/50 |
| PE/Cyanine7 Mouse IgG1, $\kappa$ Isotype | BioLegend | 400125 | 1/200 |
| Alexa Fluor® 700 Mouse IgG1, $\kappa$ Isotype | BioLegend | 400143 | 1/100 |
| Alexa Fluor® 488 Mouse IgG2a, $\kappa$ Isotype | BioLegend | 400233 | 1/100 |
| Human Phospho-STAT3 (Y705) Antibody | R&D systems | AF4607 | 1/200 |
| Systems Human Phospho-STAT5a/b | R&D systems | MAB41901 | 1/200 |
| 7-AAD Viability Staining Solution | BioLegend | 420404 | 1/100 |

**Table 6.**
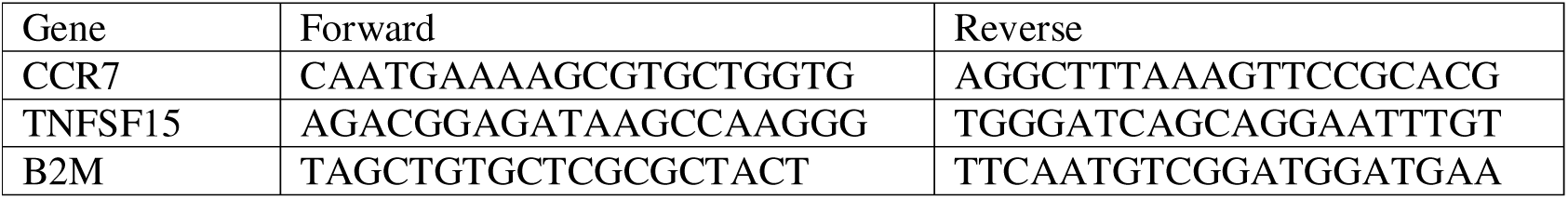
Primer sequences.

### Flow cytometry analysis

Immunophenotyping and FACS were performed on primary cells and differentiated cultures. First cells were incubated inhuman FcR Blocking Reagent (Miltenyi Biotec, #130-059-901), 5min at RT, then resuspended in 200µl staining buffer and incubated with antibodies (see Table 5) for 20 mins in the dark at 4°C. Cells were washed with cold staining buffer 3 times and filtered through 70µm filters. Cells were sorted into collection buffer consisting of 50% FBS (Thermofisher, #10099141) in HBSS (Thermofisher #14175103) using a BD FACS ARIA III/ FACS DiVa version 9 software (Becton Dickinson, Franklin Lakes, NJ) fitted with a 100um nozzle at 20psi. For immunophenotyping, CytoFLEX LX/ CytExpert software (Beckman Coulter, Brea, CA) was used to acquire samples. Post-acquisition analysis was performed using FCS Express v7 software (De Novo/ Dotmatics, Boston, MA).

### Stimulation assay

Differentiated CD11c+ CD1c+ cells from iPSC cell lines and CB donors were FACS sorted (**Table 5**) and stimulated with a combination of 1 µg/ml LPS (InvivoGen, #tlrl-smlps), 5 µg/ml HMW poly(I:C) (InvivoGen, #581942011) and 5 µg/ml R848 (InvivoGen, #tlrl-r848) in the 24 well tissue culture plate. After 16 hr, cells were collected and analysed by flow cytometry for the expression of co-stimulatory molecules (HLA-DR, CD40, CD80, CD86) (**Table 5**). The supernatant was collected for the cytokine bead assay (CBA) and was processed at the Hudson Institute, Monash University. The fluorescent intensity data for each cytokine was received, and an associated standard curve was generated. A multi-parameter regression model was fit to the standard curve and was used to calculate the concentration (pg/ml) of each cytokine.

### Real time qPCR assay

mRNA extraction of samples was carried out using Qiagen RNeasy® Plus micro Kit (Qiagen, cat#74034) and was transcribed into cDNA using Fast SYBR Green Master Mix (Thermo Fisher Scientific, #4385612), according to the manufacturer instructions. RT-qPCR was performed using the ViiA7 QRT-PCR machine (Applied Biosystems). Relative gene expression was calculated using ΔCT method normalised to human B2M housekeeping gene (See Table 5).

### RNA-seq experiment

Human iPSC-derived cDC1 and cDC2 subsets from two iPSC cell lines and CB-derived subsets from 3 healthy donors were FACS sorted as Live HLA-DR+ CD14− CD141+ CLEC9A+ cDC1, Live HLA-DR+ CD14− CD11c+ CD1c+ DCA and Live HLA-DR+ CD14+ CD11c+ CD1c+ DCB subsets. Minimum 200×10^3^ cells were collected for each sample and lysed in lysis buffer provided by Qiagen RNeasy® Plus micro Kit (Qiagen, cat#74034) proceeded by extracting RNA using the manufacturer instructions. RNA quality (determined by RNA Integrity Number (RIN^e^) and quantity were determined by High Sensitivity RNA ScreenTape (Agilent) using Agilent 2200 Technologies Tapestation System. RNA samples were processed as described previously (33) by MHTP Medical Genomics Facility at the Monash Health Translation Precinct. Illumina NSQ2000 was used for sequencing.

### RNAseq data analysis

FASTQ files were obtained from the sequencing facility and were processed using the standard Stemformatics data processing pipeline (34). Raw counts were analysed using Limma package (35). Low count genes were filtered using the default settings of filterByExpr function from EdgeR (36) while counting for the variance between biological groups. Filtered counts were normalized to logCPM and used for further analysis. A voom-plot was used to assess the quality of filtering. Clustering of samples was done using plotMDS. Ggplot2 (37) was used to generate violin and box plots. Bulk mRNA-sequencing data is available through accession GSE26173.

### Differentially expressed (DE) gene analysis

Statistical DE analysis was performed using R. The differential gene expression analysis between two groups of samples coming from different batches was performed using linear mixed-effect models by the lmer function from the lme4 package (38). In this modelling, the variable of interest and the batch parameter were considered as random effects. Benjamini-Hochberg (BH) adjustment was used for the p-value correction method. The genes with p-adjust<0.05 and logFC>1 were selected as DE genes. Code is available from the Wells laboratory github https://github.com/wellslab

### Projection method

The projection of RNA-seq data to the reference Stemformatics DC atlas has been described previously (34). The projection vignette is provided on the Stemformatics.org atlas website.

## Results

### Establishing an EB-based iPSC differentiation method to generate human DC subsets

We first set out to assess two approaches for *in vitro* DC generation, including one previously reported from human iPSC by Sachamitr *et al.* (39) and one from human cord blood (CB) CD34+ progenitors by Balan et al. (17), here referred to as Sachamitr iPSC and Balan CB methods, respectively. The Sachamitr iPSC differentiation method required a 2-week-long culture to generate hematopoietic progenitors (iPSC-HCS) from embryoid bodies (EB) in a growth factor cocktail that included GM-CSF from the beginning and then a further 10 days to differentiate these progenitors to iPSC-DC using GM-CSF and IL4 (Figure 1A). The Balan CB method included a 7-day amplification step of HSCs, then differentiation in a panel of FLT3-L, GM-CSF and IL-4 to derive DCs (Figure 1B).

**Figure 1.**
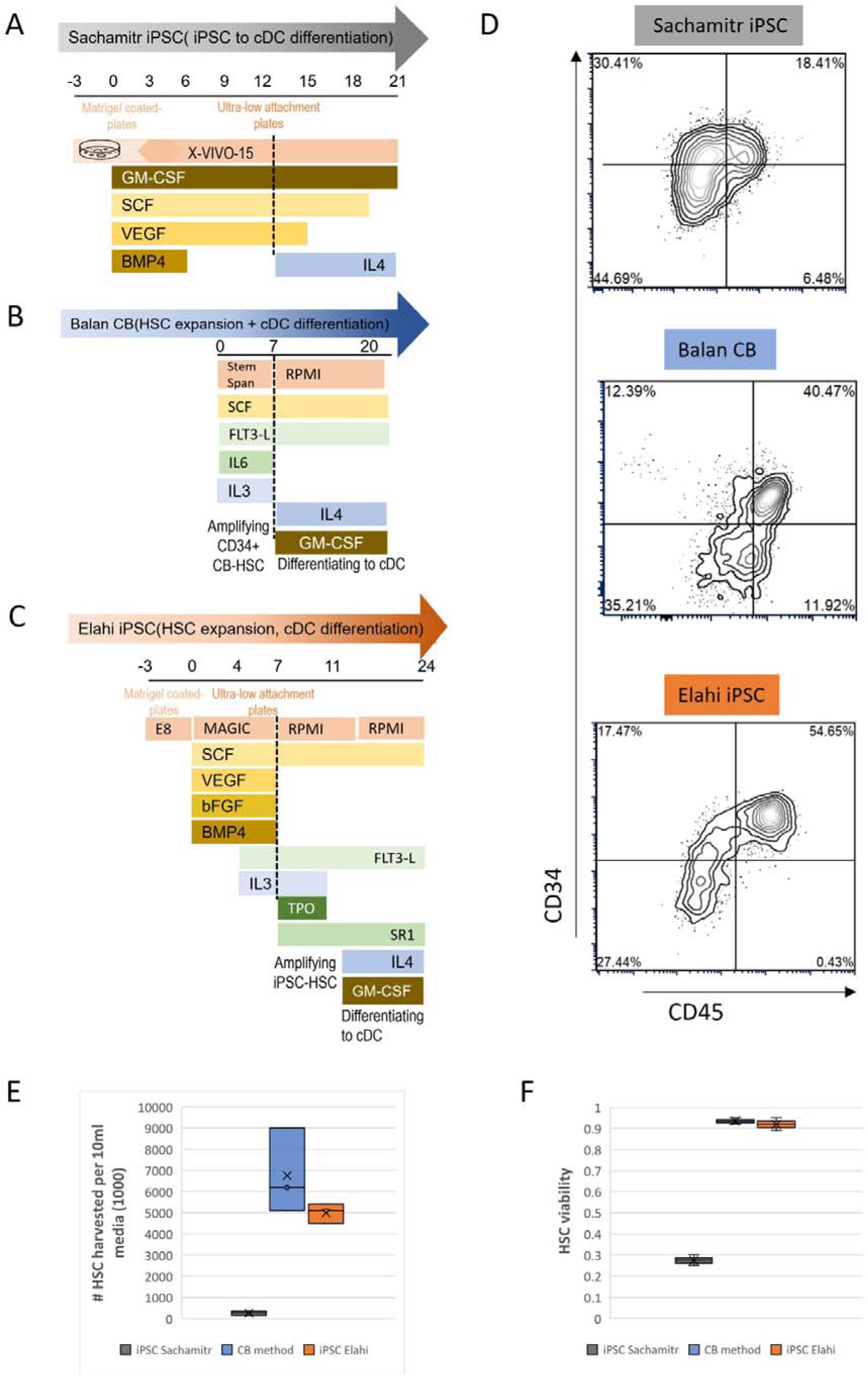
Generating hematopoietic progenitors of dendritic cells in vitro. **A)** Schematic overview of a replicated iPSC-DC differentiation method developed by Sachamitr et al. (39). **B)** differentiation protocol of cord blood (CB)-progenitors toward DC developed by Balan (17) with minor modifications. **C)** our new developed differentiation protocol of iPSC toward DC (iPSC Elahi method). **D)** Flow cytometric assessment of harvested hematopoietic stem cells (HSCs) from each method for the expression of CD34 and CD45 markers. The **E)** Number and **F)** viability of harvested hematopoietic stem cells (HSCs) from each method per batch of 10ml media. Data are representative of three replicates of two iPSC cell lines (PB001.1 and HDF51) and three individual cord blood donors (summarised at Supp. Table 2).

We designed a new protocol, referred to as the iPSC Elahi method, which combines a modified version of a contemporary EB method (28) with an amplification step of the HSC progenitors inspired by Balan *et al.* (17) (Figure 1C). Our modified four-step protocol used a brief pulse of CHIR to induce mesodermal differentiation in iPSC, then patterned embryoid bodies with BMP4, SCF, and VEGF before introducing FLT3L and IL3 to promote the production of CD34+ CD45+ HSC progenitors. We collected iPSC-derived HSC cells on day 7 of EB formation, and expanded these in FLT3L, TPO, IL3 and SR1 for 4 days (Figure 1C) to increase the number of iPSC-HSC and dendritic cell progenitors. Dendritic cells were then differentiated in media containing FLT3L, SR1, GM-CSF and IL4 for up to two weeks.

The Sachamitr iPSC protocol did generate ∼18% CD34+ CD45+ cells at day 13, indicating specification to a hematopoietic cell type, but with low numbers of HSCs harvested (<300×10^3^ cells per 10ml batch) with 30% viability (Figure 1 D-F). In contrast, the Balan CB method showed a 40% CD34+ CD45+ HSC yield, generating >5×10^6^ progenitors with 95% viability (Figure 1 D-F). Despite implementing multiple levels of improvement to the iPSC Sachamitr protocol, the quantity of generated HSCs by this method still fell significantly short of those produced by the CB method (described in Supp. Method, Supp. Table 1 and Supp. Figure 1). We were motivated to develop a new iPSC protocol that generates a similar or even higher proportion of HSC progenitors than the CB method. Our method (Elahi iPSC) significantly enhanced the yield of the CD34+ CD45+ HSC generation by up to 54% and increased their number to more than 10-fold compared to the previous Sachamitr iPSC method (Figure 1 D-F). However, the number and the viability of generated HSC were similar between our method and the CB differentiation approach.

Assessing the terminally differentiated DC from our expanded iPSC-HSC, we observed that 2×10^6^ of iPSC-HSC yielded 10-12% cDC1-like cells expressing CLEC9A+ CD141+, of which we routinely generated 1.8×10^3^ DC1 per batch of 10ml media, and an equivalent number of cDC2-like cells expressing CD11c+ CD1c+ (Figure 2). This was more than the number of cDC2 and equivalent to cDC1 obtained from the Balan CB method in our hands. The proportion of cDC-like subsets obtained from the Sachamitr protocol was significantly lower than that of our protocol. The high number of DCs obtained by the Elahi iPSC method was greatly enhanced by increasing the number of progenitors available for differentiation. From this point forward, we will stop characterising the Sachamitr method, and any dendritic cells derived from Elahi iPSC method will now be referred to as iPSC-DC.

**Figure 2.**
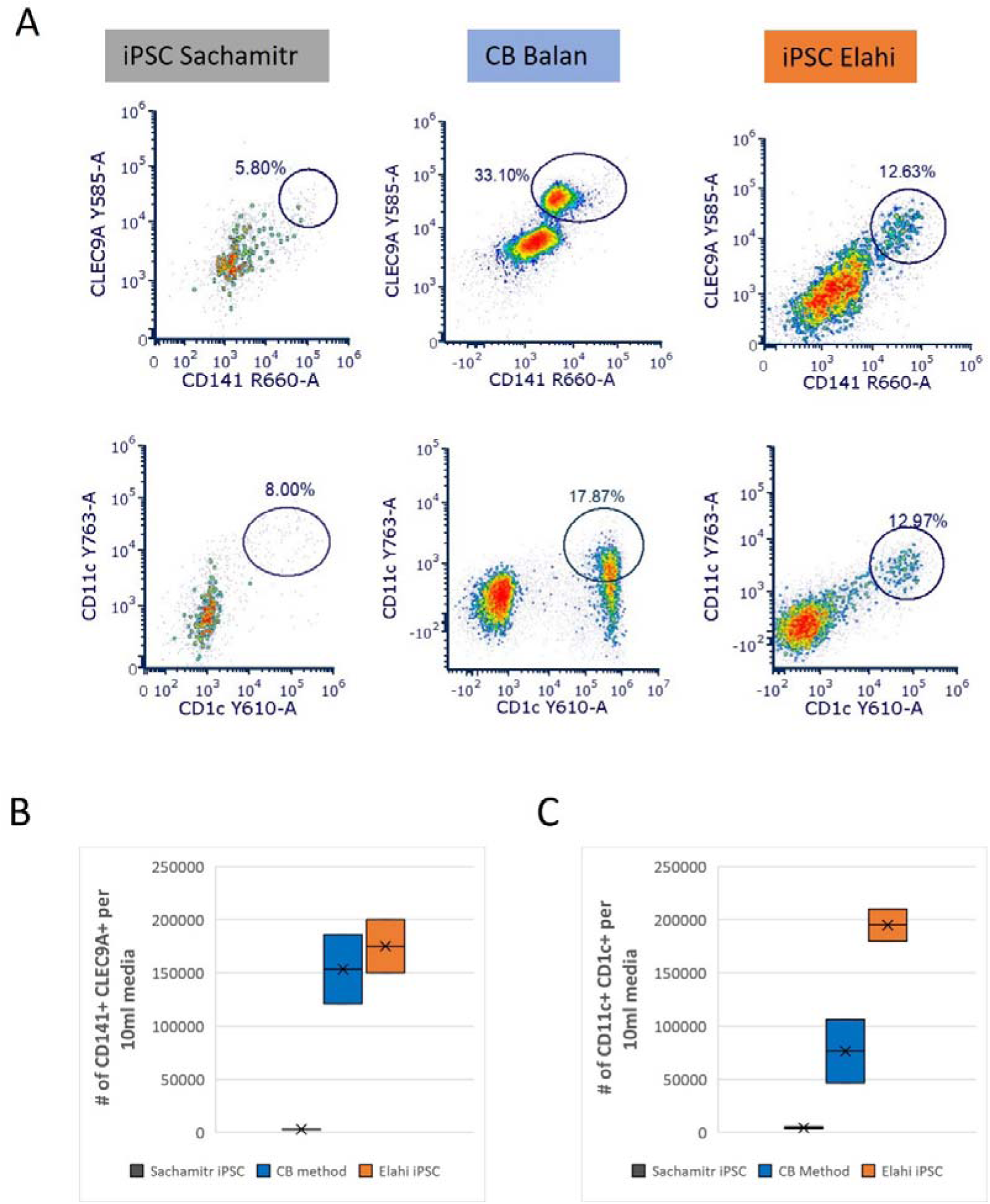
New developed protocol is more efficient in producing iPSC-derived conventional dendritic cells. **A)** The flow cytometry analysis of terminal differentiated cDCs comparing the percentage of the differentiated Live CD141+ CLEC9A+ cDC1 (top) and Live CD11c+ CD1c+ cDC2 (bottom) between the replicated Sachamitr protocol, cord blood (CB) method and our new developed protocol (Elahi iPSC). **B)** Box plot showing the number of differentiated CD141+ CLEC9A+ cDC1 and **C)** CD11c+ CD1c+ cDC2 per 10ml media compared across multiple protocols including Sachamitr protocol, cord blood (CB) method and our new developed protocol (Elahi iPSC). Data is from two replicates of PB001.1 iPSC cell line and two replicates of one CB donor.

### CD11c^+^ cells differentiated from iPSC *in vitro* cannot be efficiently generated by FLT3L alone

We next sought to optimise the number of FLT3L+ CD11c+ DC. Four conditions were tested using two different iPSC lines (HDF51 and PB001.1). FL: 100ng/ml FLT3L alone; FLG: FL+ 20ng/ml GM-CSF; FLI: FL+20ng/ml IL4 or FLGI which combined all three cytokines. FLT3L alone did not efficiently generate CD11c+ cells (Figure 3), however, in combination with GM-CSF, gave rise to 5% CD11c+ cells. The most efficient combination was FLGI culture, which gave rise to 24.9% CD135+ CD11c+ cells, showing that the combination of GM-CSF and IL-4 is essential for the development of the iPSC-DC. We then sought to find the optimum concentration of these cytokines. Increasing the amount of GM-CSF/IL4 relative to FLT3L from 5ng/ml to 50 ng/ml promoted the generation of CD14+ cells, whilst the proportion of CD141+ cells remained the same. We, therefore, set the optimum concentration of GM-CSF and IL4 for iPSC-DC differentiation at 5 ng/ml.

**Figure 3.**
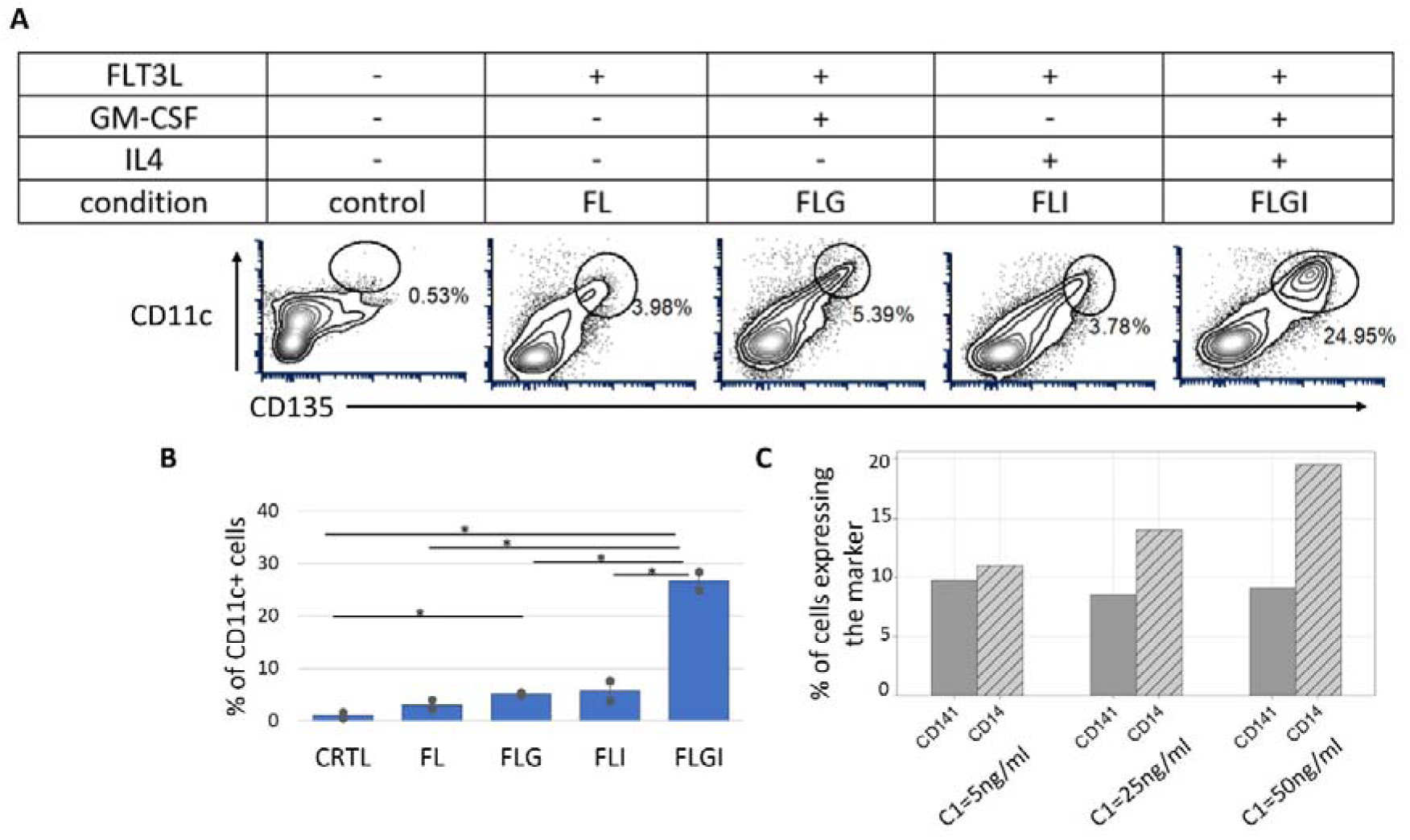
The in vitro differentiation of CD135+ CD11c+ cells require a combination of GM-CSF and IL4 cytokines on top of the FLT3 ligand. **A)** Representative flowcytometric plots assessing the yield of CD11c+ generation under four conditions including FL (100ng/ml FLT3L), FLG (100ng/ml FLT3L+ 20ng/ml GMS-CF), FLI (100ng/ml FLT3L+ 20ng/ml IL-4), FLGI (100ng/ml FLTL3+ 20ng/ml GM-CSF+ 20ng/ml IL-4) and control (no FLT3/GM-CSG/IL4). B) The bar graph compares the percentage of CD11+ cells under the four conditions mentioned earlier. Data were obtained from two iPSC cell lines, PB001.1 and HDF51. (*P<0.05, student T-test). C) The grouped bar graph showing the percentage of CD141+ and CD14+ cells differentiated under three different concentrations of GM-CSF/IL4, including C1:5ng/ml, C2:25ng/ml and C3:50ng/ml. In all conditions, FLT3L is present at 100ng/ml.

### iPSC-cDC subsets derived *in vitro* are phenotypically heterogeneous

To investigate the identity of the dendritic cells derived from iPSC *in vitro*, we ran a flow cytometric analysis on cells at day 23 of differentiation. We used a panel of antibodies, including HLA-DR, CD11c, CD141, CLEC9A, CD1c and CD14, in addition to the viability stain. To combat highly auto-fluorescent (AF) cells, we introduced gating steps in short and long wavelengths to remove them (Figure 4 A). The iPSC-derived cDC1-like subset was characterised as Live HLA-DR^+^ CD14−CD11c^mid/+^ CD141+ CLEC9A+.

**Figure 4.**
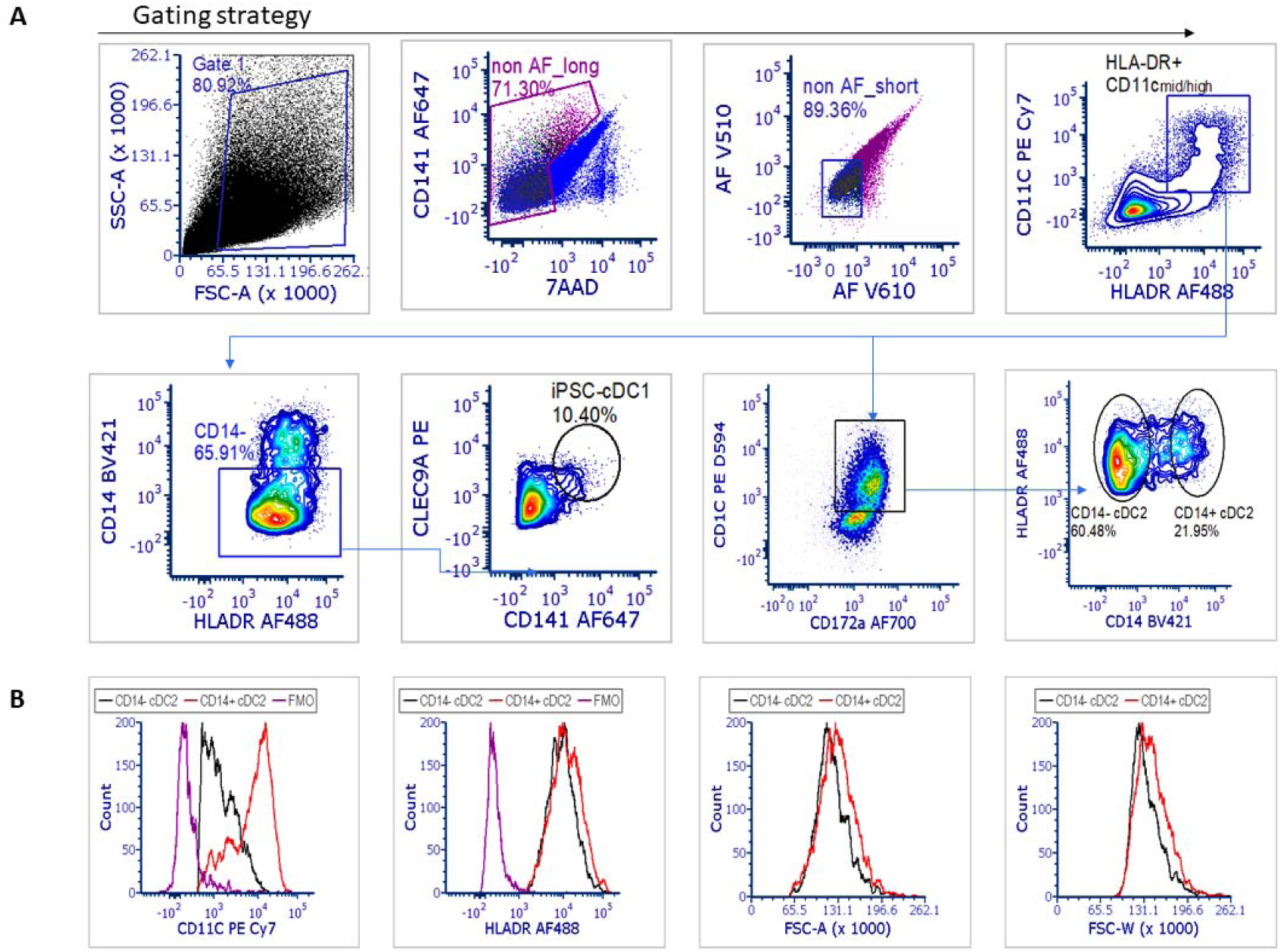
Flow cytometric characterisation of differentiated iPSC-cDC subsets. A) Auto fluorescent cells (AF) were removed from further analysis during two steps. iPSC-cDC1 and iPSC-cDC2 were characterized as Live HLA-DR^+^ CD14− CD11c^mid/+^ CD141+ CLEC9A+ and Live HLA-DR^+^ CD11c+ CD172a+ CD1C+ cells, respectively. B) Assessment of iPSC-derived CD14+ cDC2 and CD14− cDC2 subpopulation regarding their differences in CD11c and HLA-DR expression levels. The area and width of the forward scatter signal (FSC-A and FSC-W) represent their size and granularity, respectively. CD14+ population is in black, CD14− in red and the FMO (full minus one) control in black.

The iPSC-derived cDC2-like subset was characterised as Live HLA-DR^+^ CD11c+ CD172a+ CD1C+ cells. This population was heterogeneous in terms of the expression of CD14, including CD1c+ CD14− and CD1c+ CD14+ groups (Figure 4 A). CD1c+ CD14+ cDC2 showed a greater level of CD11c expression than CD1c+ CD14− cDC2, while the level of HLA-DR expression was similar (Figure 4 B). The CD14+ population was slightly larger by having a higher area of the forward scatter signal (FSC-A). They also had marginally more granularity than CD14− cDC2, measured by the width of the forward scatter signal (FSC-W).

### iPSC-derived dendritic cells (iPSC-DCs) are not transcriptionally equivalent to primary cDCs

To understand the biology of iPSC-derived DC (iPSC-DC) subsets and to investigate their similarity with cord blood-derived DC (CB-DC) equivalents, we FACS sorted each subset and analysed them through RNA sequencing (Figure 5 A). Both CB-derived DC and iPSC-DC were expanded at the progenitor stage for 4 days before differentiation. The DC1 subset was sorted as Live HLA-DR^+^ CD14− CD11c^mid/+^ CD141+ CLEC9A+. Two populations of DC2 were sorted as Live HLA-DR^+^ CD11c+ CD172a+ CD1C+ with negative or positive expression of CD14 referred to as DC2A and DC2B, respectively. At least 200×10^3^ cells were sorted for each sample with replicates from two different iPSC cell lines and from three CB donors, noting that the cDC1 population from one CB donor failed RNA QC because of low cell numbers.

**Figure 5.**
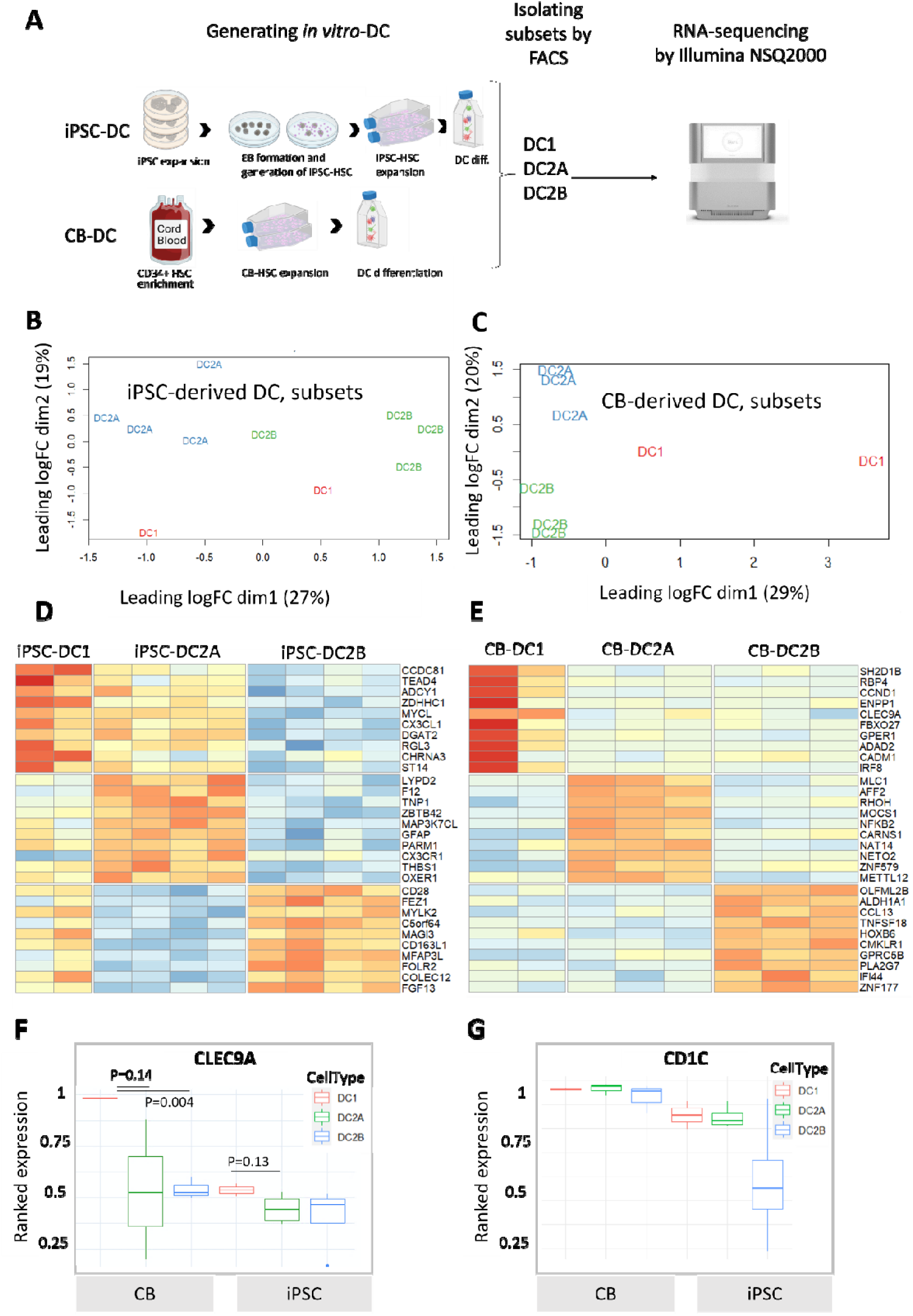
Characterisation of iPSC- and CB-derived dendritic cells by bulk RNA-sequencing. A) Experimental setup of generation DCs from iPSC and cord blood (CB). The process includes iPSC expansion, harvesting of haematopoietic stem cells (iPSC-HSC), expansion of iPSC-HSC, and differentiating to dendritic cells. Similarly, CB-HSC were expanded and differentiated to DCs. Samples were FACS sorted and analysed by bulk RNAseq. B) Multidimensional scaling (MDS) plot of log2 fold changes showing sample clustering of iPSC-derived dendritic cells (iPSC-DC). C) Multidimensional scaling (MDS) plot of log2 fold changes showing sample clustering of cord blood CD34+-derived DC (CB-DC). D) Heatmap of differentially expressed (DE) genes by iPSC-DC subsets. The top 10 DE genes with p_adjust (BH)<0.01 ranked with logFC were selected for heatmap display (See extended list in Supp. Table 3). E) Heatmap of differentially expressed (DE) genes by CB-DC subsets. The top 10 DE genes with p_adjust (BH)<0.01 ranked with logFC were selected for heatmap display (See extended list in Supp. Table 4). F) CLEC9A ranked gene expression among iPSC-DC and CB-DC subsets. G) CD1C ranked gene expression among iPSC-DC and CB-DC subsets. P-value calculated by student T-test.

We first examined the sample clustering using a multidimensional scaling (MDS) plot. The first dimension (dim 1) accounted for 27% of expression variability and defined subsets of iPSC-DC2A and iPSC-DC2B, respectively. The second dimension (dim2) accounted for 19% of expression variability and described the difference between iPSC-DC1 and iPSC-DC2s (Figure 5). In contrast, the CB-cDC1 and cDC2 subsets were captured on dim 1, 29% of expression variability, while dim 2 (20% variability) separated CD14+ and CD14− cells (Figure 5 C).

Next, we identified the differentially expressed genes that separated subsets within the iPSC-DC groups (Supp. Table 3). The 10 most discriminating genes ranked by LogFC were chosen for illustrative purposes (Figure 5 D). The genes highly expressed by iPSC-DC1 were closely related to the maturation and migration of dendritic cells (e.g. MYCL, CXCL1, ZDHHC1, SEMA3C, NR4A3) and the antigen presentation process (e.g. TEAD4, ERAP2, AP1S1). iPSC-DC2A upregulated genes involved in communication with T cells (e.g. THBS1, FCER1A, CD1D, CD19) and iPSC-DC2B showed upregulation of cytokine signaling genes (e.g. CD28, IL10, TNFSF12, IL15RA). We asked whether genes that exhibit distinct expression patterns in iPSC-DC subsets also demonstrate subset-specificity across DC subsets derived *in vivo*. Our analysis, leveraging the Human DC Atlas database (40), revealed that most of these genes do not display subset-discriminating patterns among *in vivo* DCs (Supp. Figure 2A).

Investigating the differential expression profile of the CB-derived DCs showed high expression of CLEC9A, IRF8, CADM1 and XCR1 by CB-DC1 subset (Figure 5 E, Supp. Table 4). This profile suggests that CB-DC1 successfully captured the identity of the DC1 subset. The DC1 identity was partially obtained by iPSC-DC1 as they exhibit higher levels of CLEC9A and CADM1 compared to iPSC-DC2A/B but lack the expression of IRF8 and TLR3 (Supp. Figure 2B). However, the level of CLEC9A expression by CB-DC1 remain notably higher compared to iPSC-DC1 (Figure 5 F). CB- and iPSC-derived DC1 cells appear to have a hybrid transcriptional profile by expressing high levels of the CD1C gene, which is a common marker of the cDC2 subset (Figure 5 G). Altogether, iPSC-derived DCs captured a transcriptional identity that is different from their primary DC equivalents.

We then examined the projection of the sequenced samples to the reference Human DC Atlas. The iPSC-DC subsets strongly clustered and showed transcriptional similarity with the *in vitro* (Mo-DC) samples (Figure 6 A-B). To determine whether iPSC-DC, with a monocytic profile, are distinct from the iPSC-derived macrophages (ipMAC), we combined our dataset with another RNAseq dataset from our laboratory (28) that has profiled ipMAC and macrophages differentiated from blood monocytes (MDM). To be consistent with our cells, the samples selected from this dataset were non-activated control MDM and ipMAC samples. Previously, IRF4 and MAFB transcription factors were discovered to distinguish human MoDC from MoMAC and vice versa (41). Consistent with this finding, our ipDCs showed higher expression of IRF4 but lower MAFB compared to ipMAC and MDM (Figure 6 C-D).

**Figure 6.**
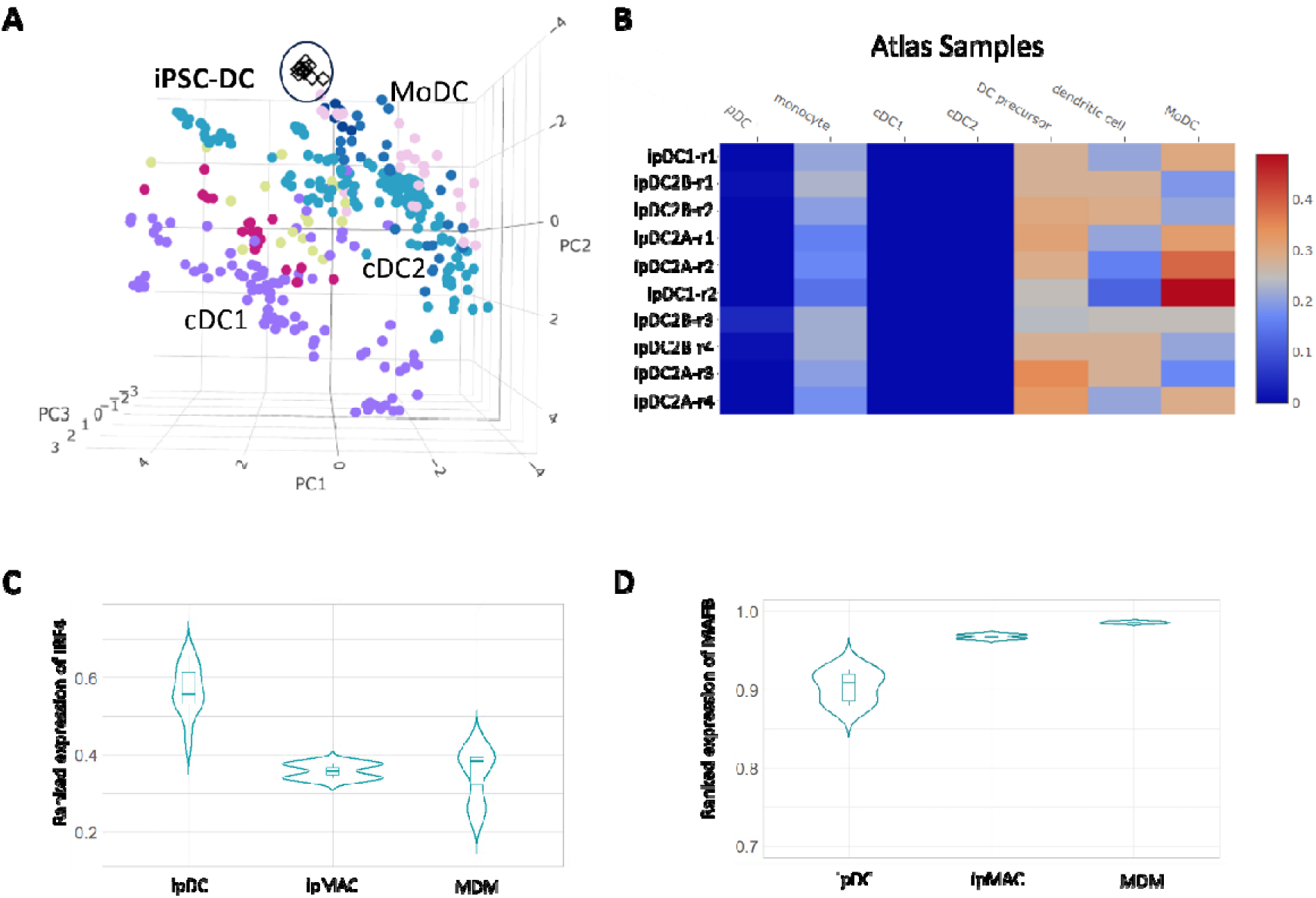
Benchmarking iPSC-derived DCs against other myeloid cells. A) Projection of iPSC-DC on Human DC Atlas showing the proximity of iPSC-DC transcription with atlas MoDC subset. B) Capybara similarity scores between iPSC-DC and atlas cell types showing that iPSC-DC capture the MoDC transcriptional profile. C) Violin plot of ranked gene expression of IRF4, and D)MAFB transcription factors compared between our iPSC-derived DCs (ipDC) with external transcriptional data (28) on iPSC-derived macrophages (ipMAC) and macrophages differentiated from blood monocytes (MDM), previously obtained in our laboratory.

### iPSC-derived DCs respond to TLR stimulation as efficiently as CB-DC2s

To examine the immunoactivation capacity of iPSC-DC and CB-DC, we selected a combination of LPS, Poly I:C and R848 (referred to as LPR stimuli) which are commonly used to test the DCs maturation. FACS-sorted CD11c+ CD1c+ iPSC-DC and Cd11c+ CD1c+ CD14− CB-DC2A were stimulated overnight (16 hr) and assessed for secreted immune modulators by cytokine bead assay (CBA). Upon stimulation, both iPSC-DC and CB-DC2 secreted pro-inflammatory cytokines, including TNFα, IL-6 and IL-8 in high levels and IL-1 (IL-1a and IL-1b) in lower levels (Figure 7). This highlighted their similarity with the known functional characteristics of cDC2s. Interestingly, non-stimulated (control) CB-DC2 samples were able to secrete a moderate level of TNFα, IL-8 and IL-1a in contrast to control iPSC-DC with no response.

**Figure 7.**
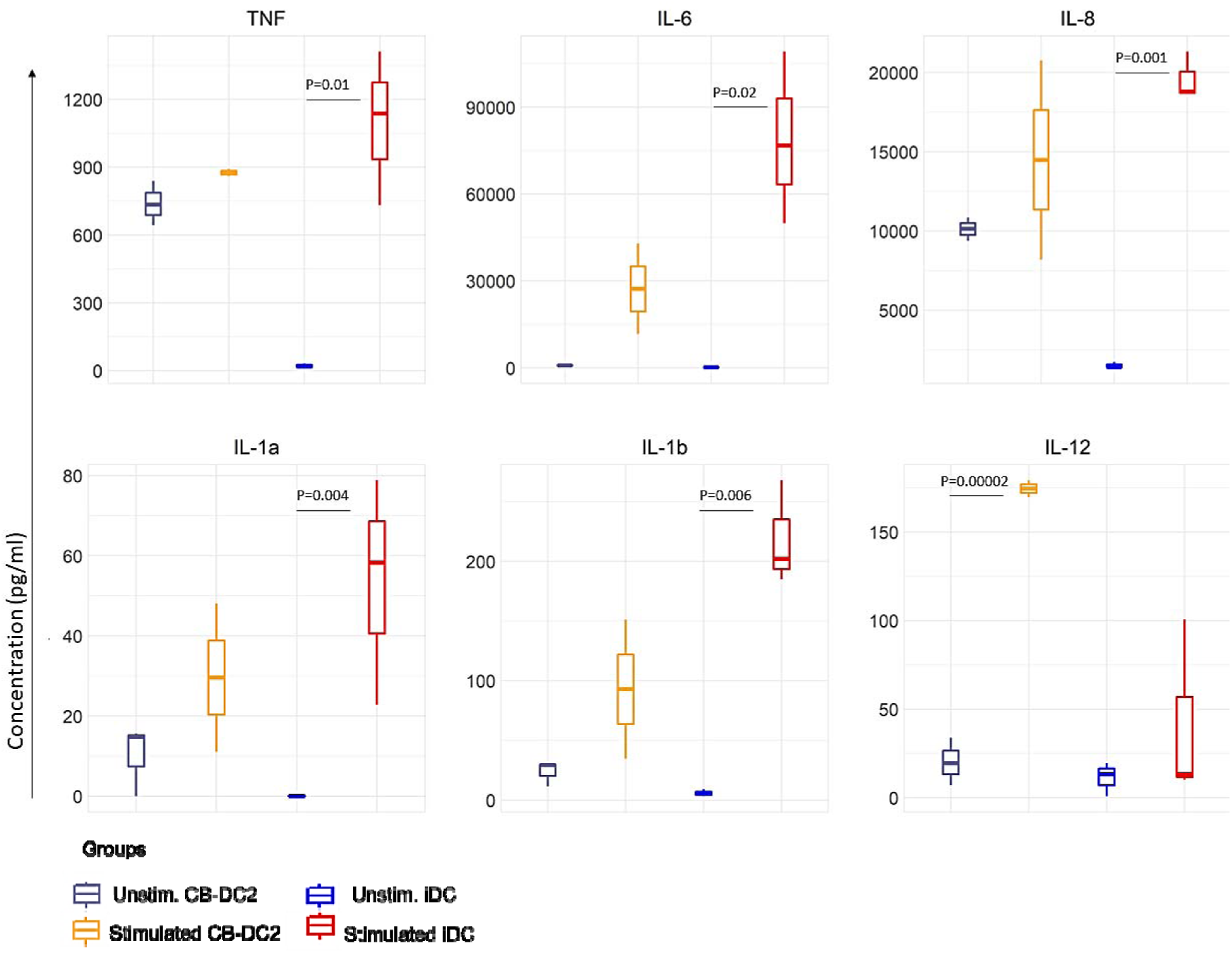
iPSC-DC are activated by TLR adjuvants as efficiently as CB-DC2A. FACS-sorted CD1c+ iPSC-DC and CD1c+ CD14− CB-DC2A were stimulated overnight (16 hr) and assessed for secreted immune modulators by cytokine bead assay (CBA). Three samples of iPSC-DCs are differentiated from HDF51 and PB001.1 lines. Three samples of CB-DC differentiated from CD34+ cord blood cells of two donors. P-value calculated by student T-test.

Furthermore, the ability of iPSC-DC to enhance the expression of co-stimulatory molecules and upregulation of CCR7 in response to LPR stimuli was assessed. iPSC-DC exhibited a robust upregulation of CD80 and CD40 following activation, with a median fluorescent intensity (MFI) approximately double that of unstimulated samples (Figure 8). However, a slight upregulation of CD86 and HLA-DR was observed; the differences between stimulated and unstimulated samples were not statistically significant. We also observed a significant upregulation of the migration-associated gene, CCR7, and the inflammatory TNFSF15 gene by iPSC-DC after stimulation compared to the control unstimulated samples (Supp. Figure 3). To conclude, the functional analysis confirmed that the in vitro-derived DCs generated from iPSC by our method are able to respond to the TLR stimuli.

**Figure 8.**
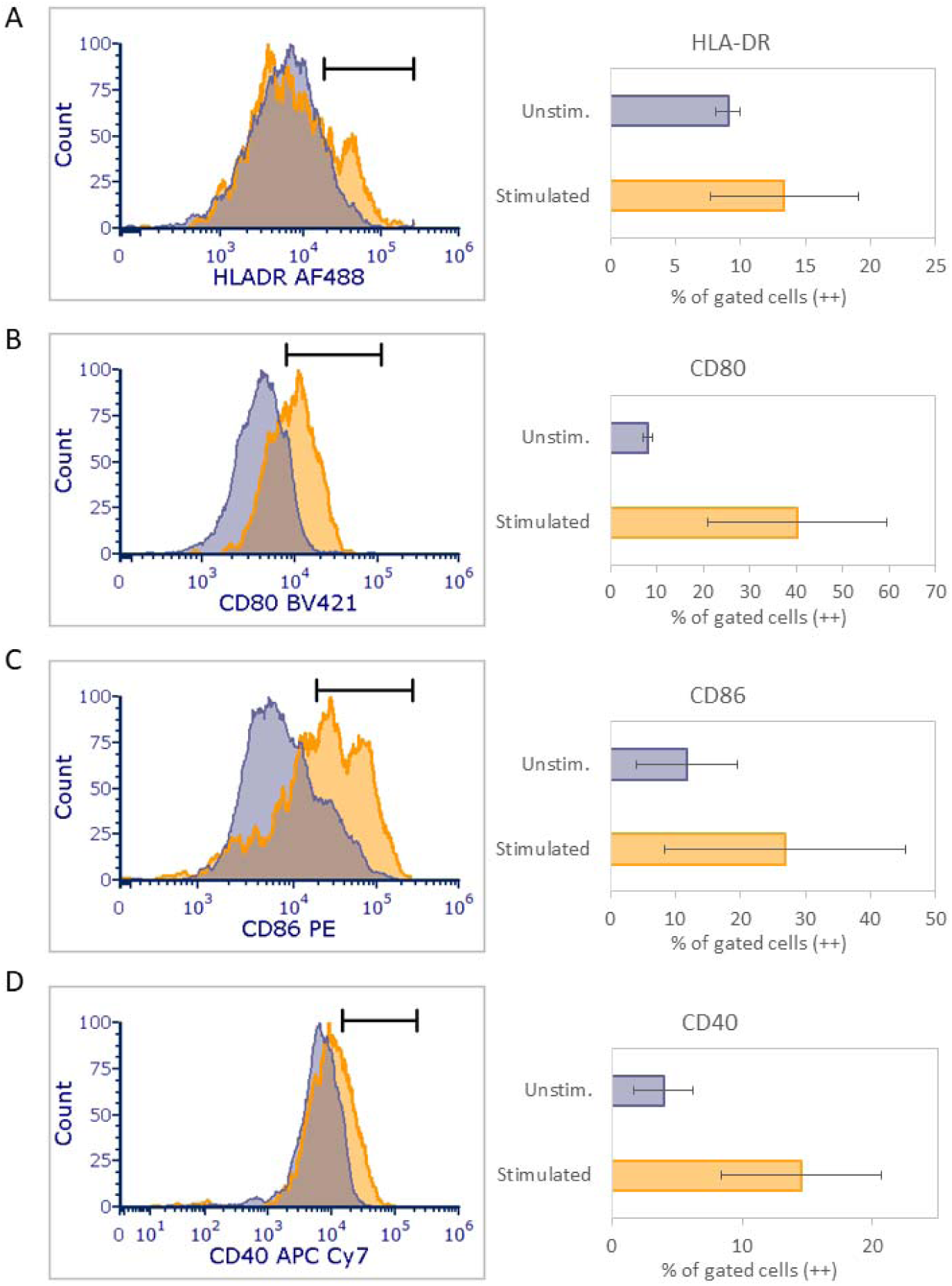
Enhancement of co-stimulatory molecules’ expression after activation of iPSC-DC. Analysis of median fluorescent intensity (MFI) of A) HLA-DR and co-stimulatory molecules B) CD80, C) CD86 and D) CD40 between stimulated and unstimulated FACS sorted CD1c+ iPSC-DC in blue and grey, respectively. For each condition, four samples including two replicates of HDF51 and PB001.1 cell lines, were tested. P-value calculated by student T-test.

## Discussion

We designed and developed an optimised protocol for generating human dendritic cells from iPSC in a feeder-free differentiation approach. Our method incorporates an amplification phase of iPSC-derived myeloid progenitors, resulting in an improved differentiation yield and a greater number of iPSC-derived DCs compared to a previous iPSC method and similar to the yields obtained from cord blood *ex vivo* expansion methods. Given the limitation in CB methods is the amount of available starting materials, the iPSC method described here offers a promising approach for scale up of human DC production.

The terminally differentiated iPSC-derived DC by our method expressed the markers of cDC subsets at the protein level; however, the mRNA expression profile of these cells showed a hybrid phenotype that was most similar to moDC. This inconsistency between gene and protein expression in our iPSC-DC subsets may be due to post-transcriptional regulation. Protein level depends not solely on mRNA levels, but also on translation rate and protein half-lives (reviewed by Buccitelli and Selbach (42)). Therefore, the imperfect protein-mRNA correlation can arise from translation efficiency and protein half-live differences. Non-proliferative cells, a behaviour that we observed in our iPSC-DCs, have protein half-lives greater than 500 hours (43), which may result in a stable protein level linked with lower mRNA levels.

Our transcriptional analyses of iPSC-derived DC subsets further highlights that these cell types have a hybrid phenotype, which is distinct from primary DCs developed *in vivo*. For example, iPSC-DC1 show the expression of DC1-specific genes (e.g., CLEC9A and CADM1) but they also expressed DC2-markers (e.g., CD1C), lacking the full identity of either subset. This hybrid identity has been previously seen in the culture of CB progenitors under FLT3L-driven differentiation. Culture of primary progenitors can induce expression of the CD1C marker in cDC1s (18, 44) and others have shown that iPSC-derived DC1 co-express XCR1 and CD14 markers (39, 45). This mixed phenotype can be attributed to the culture conditions under which the signals from the environment may not target a specific transcriptional network necessary for deriving a particular DC subset. To our knowledge, the generation of DC1 from iPSC under a feeder-free culture system has not yet been achieved. This is also consistent with previous studies that reported a MoDC/DC2-like profile (26, 46, 47) or the expression of CD14 monocytic marker by their iPSC-derived DCs (39, 45).

Given iPSC-DCs have a hybrid phenotype that at least mimic primary moDC, they have the potential for translational studies. Ultimately for clinical applications, it is important that the *in vitro*-generated DC are functional in that they are migratory to lymph nodes, can recognise pathogens, and are capable of processing and presentation of antigens to T-cells, while producing inflammatory cytokines that provide an instructive environment. Our stimulated *in vitro* iPSC-derived DCs appear to be able to produce pro-inflammatory cytokines as well as upregulate appropriate co-stimulatory and migration molecules, although in vivo migration, and engagement with T cells has not yet been tested.

## Author contributions

ZE conceptualisation, methodology, and writing the original draft. CAW Supervision, conceptualisation, writing – review and editing. VJ Formal analysis, writing – review and editing. MS Formal analysis, writing – review and editing. SKB Data curation, writing – review and editing. KR Supervision, methodology. JDM Supervision, writing – review and editing.

## Acknowledgements and funding

This work was supported by the Australian National Health and Medical Research Council APP1186371 to CAW. The authors thank the Melbourne Cytometry Platform at the University of Melbourne for infrastructure support, and the Hudson Genomics Facility, Hudson Institute of Medical Research, for assistance with RNA library preparation and sequencing.

## Supplementary Materials

To enhance the Sachamitr protocol, we firstly updated our approach to handling the iPSC, removing mechanical dissociation of iPSC in favour of an enzyme-based approach, as well as adding ROCK inhibitor at the seeding stage to improve the quality of embryoid bodies that were important for generating fresh HSCs comparable to the primary CB cells (Supp. Table 1). This modification step also enhanced the viability of harvested HSCs around EBs but did not improve the yield obtained from the Sachamitr method (Supp. Figure 1).

We next swapped XVIVO differentiation media for MAGIC media, which has been designed for haematopoietic cell differentiation. We took pains to remove the debris resulting from broken EBs, which improved the quality of culture (Supp. Figure 1), and doubled the number of HSC harvested. Our third approach to improving the quality of DC progenitors from EBs was to introduce a shaking culture system to improve the circulation of media and oxygen around the dish, plus providing a driving force for HSCs to bleed from EBs. This step also enhanced the number of harvested HSCs and their viability (Supp. Figure 1), but still failed to produce numbers that would be equivalent to those derived from cord blood.

**Supplementary Table 1.** Identified factors associated with the low quality of culture in the replicated iPSC-DC protocol and the drawbacks related to each one. For each item, a modifications approach was applied to enhance the quality and quantity of harvested iPSC-HSC.

| identified factors |  |  | Approach by replicated protocol | Drawbacks | Modification applied |
| --- | --- | --- | --- | --- | --- |
|  | 1) Cell viability at seeding step | dissociation method | mechanical, scraping | inducing cell death | enzymatic, EDTA-based (Modification 1) |
|  |  | cell survival at the seeding step | no supporting factor | low survival rate | add ROCK inhibitor (Modification 1) |
|  | 2) Culture media | culture media | XVIVO | not supporting EB growth, undisclosed components | MAGIC (based on BPEL(31) media) (Modification 2) |
|  |  | media change strategy | replacing one half of the spent media with a fresh one | leaving EB and cell debris in the culture environment | replacing whole media with new media (Modification 2) |
|  | 3) suspension culture design | static/shaking | non-shaking | less access to nutrients by EBs, low chance of bleeding from EBs for HSCs | introducing EBs to a shaking system from day 0 (Modification 3) |

**Supplementary Table 2.**
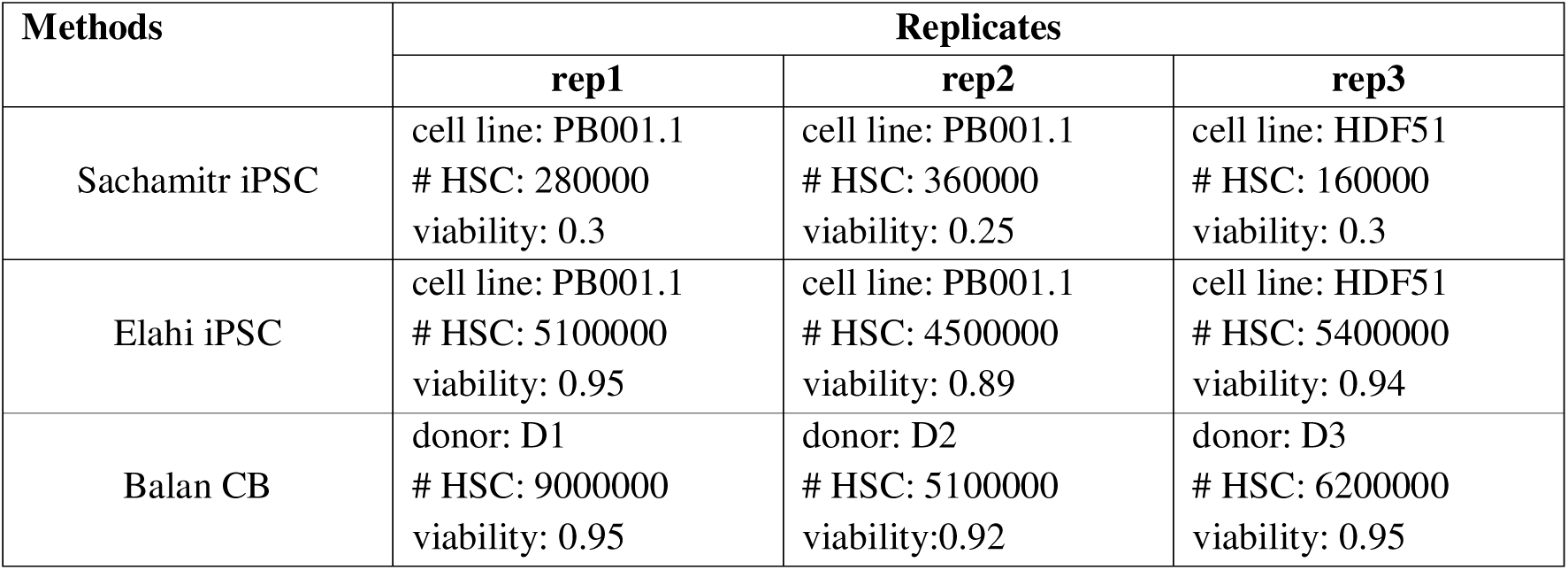
The number of hematopoietic progenitors (HSC) generated by different methods. Methods include Sachamitr iPSC, Elahi iPSC and Balan CB with the details of cell line or donor used for each experiment.

| Methods | Replicates |  |  |
| --- | --- | --- | --- |
|  | rep1 | rep2 | rep3 |
| Sachamitr iPSC | cell line: PB001.1<br># HSC: 280000<br>viability: 0.3 | cell line: PB001.1<br># HSC: 360000<br>viability: 0.25 | cell line: HDF51<br># HSC: 160000<br>viability: 0.3 |
| Elahi iPSC | cell line: PB001.1<br># HSC: 5100000<br>viability: 0.95 | cell line: PB001.1<br># HSC: 4500000<br>viability: 0.89 | cell line: HDF51<br># HSC: 5400000<br>viability: 0.94 |
| Balan CB | donor: D1<br># HSC: 9000000<br>viability: 0.95 | donor: D2<br># HSC: 5100000<br>viability: 0.92 | donor: D3<br># HSC: 6200000<br>viability: 0.95 |

**Supplementary Figure 1.**
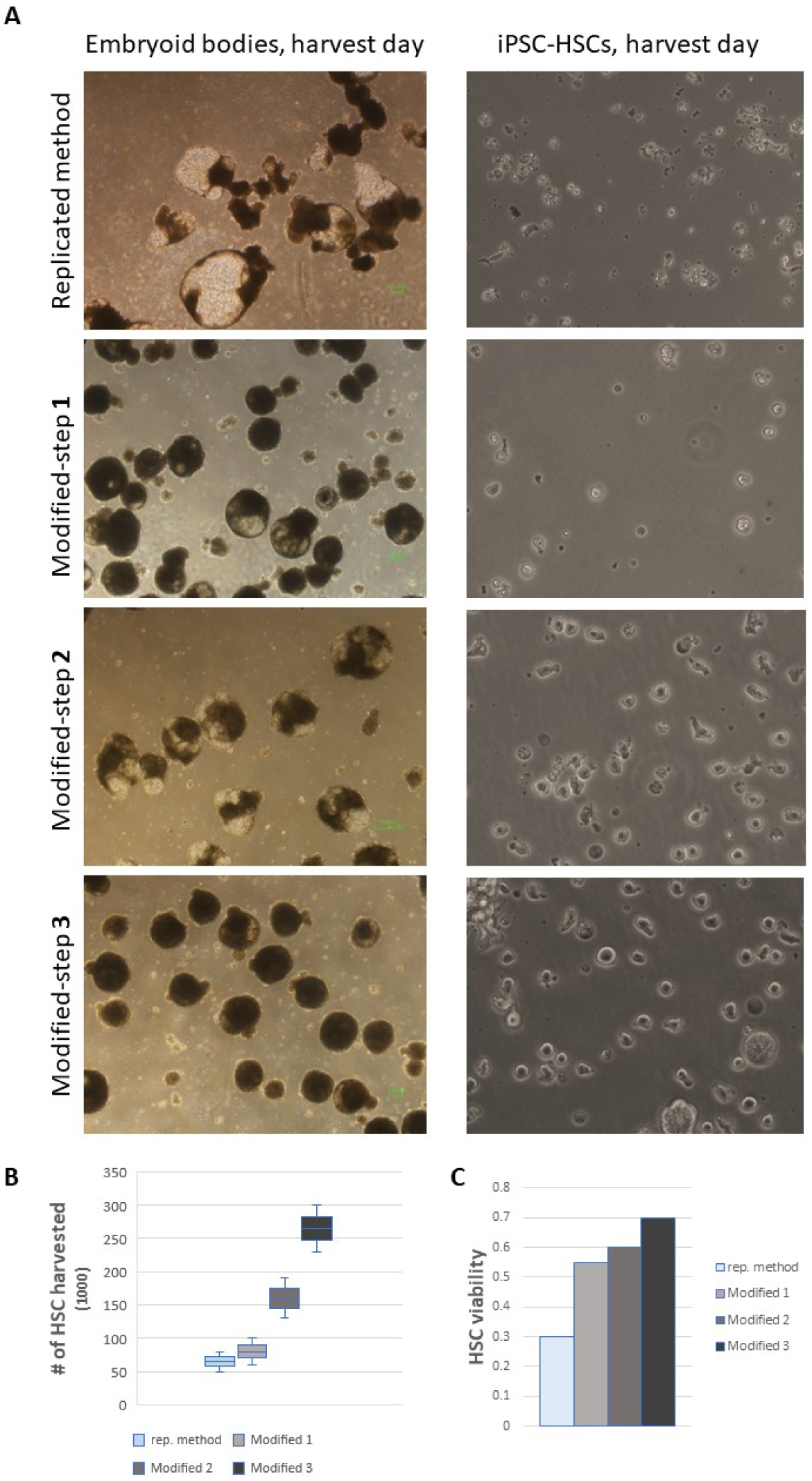
Step by step improving the quality and the number of generated HSCs from replicated Sachamitr iPSC method (Modification details are described in Supplementary Table 1). **A)** Light microscopy images of embryoid bodies (EB) at day 13 differentiation (left) and the generated HSCs around EBs (right) across each modification step of the Sachamitr protocol. **B)** Box plot showing the number of harvested HSCs and **C)** their viability at day 13 of differentiation from each modification step. Bar plot comparing the iPSC-HSC viability across multiple modification practices of the replicated protocol.

**Supplementary Figure 2.**
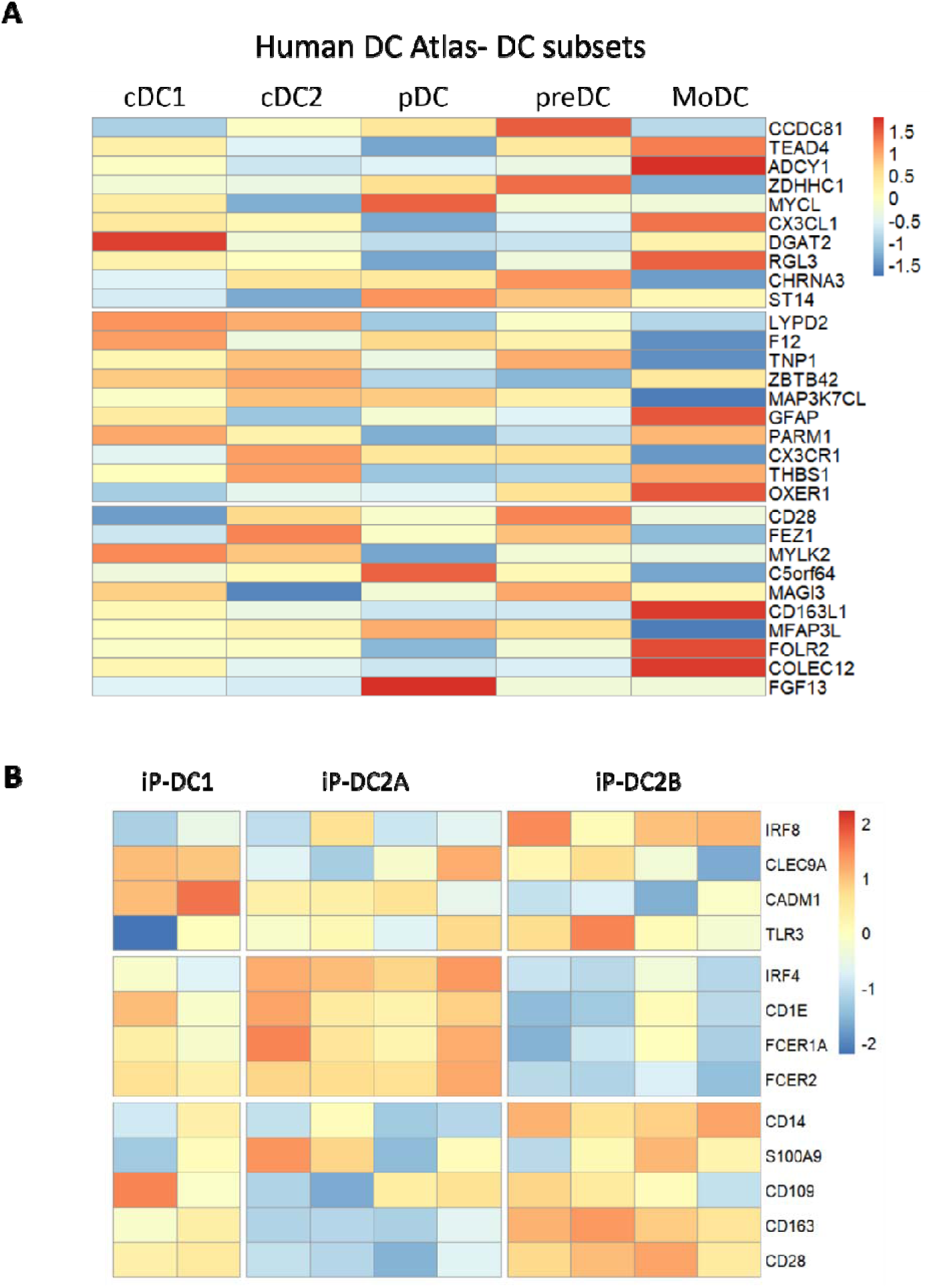
Dendritic cell subset-discriminating genes. **A)** Heatmap of the top 10 DE genes expressed by iPSC-DC subsets compared between the reference Human DC Atlas (40) cell types. The genes identified with p_adjust (BH)<0.01 and ranked with logFC among iPSC-DC subsets. The scale next to the heatmap shows the colour scale from −1.5 (lowest scaled expression in blue) to 1.5 (highest scaled expression in red). **B)** Heatmap of some known DC subset-defining genes across iPSC-derived DC subpopulations. The subset-significant genes extracted from literature. The scale next to the heatmap shows the colour scale from −2 (lowest scaled expression in blue) to 2 (highest scaled expression in red).

**Supplementary Figure 3.**
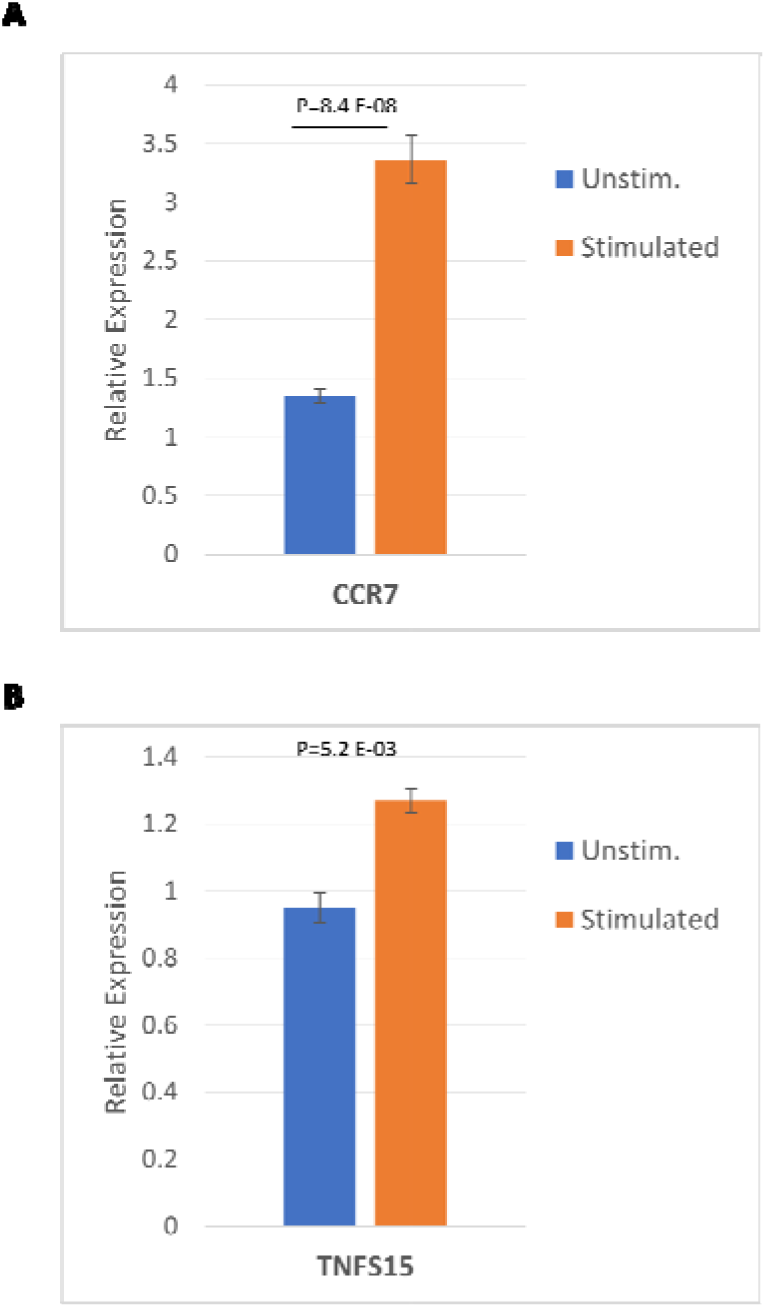
Assessing the expression of CCR7 and TNFS15 genes after activation by iPSC-DC. **A)** Boxplot showing the relative expression of the CCR7 gene associated with DC migration compared between stimulated and unstimulated conditions analysed by RT-qPCR analysis. **B)** Boxplot showing the relative expression of TNSF15 inflammatory gene compared between stimulated and unstimulated conditions. P-value: student T-test.

